# A target enrichment bait set for studying relationships among ostariophysan fishes

**DOI:** 10.1101/432583

**Authors:** Brant C. Faircloth, Fernando Alda, Kendra Hoekzema, Michael D. Burns, Claudio Oliveira, James S. Albert, Bruno F. Melo, Luz E. Ochoa, Fábio F. Roxo, Prosanta Chakrabarty, Brian L. Sidlauskas, Michael E. Alfaro

## Abstract

1. Target enrichment of conserved nuclear loci has helped reconstruct evolutionary relationships among a wide variety of species. While there are preexisting bait sets to enrich a few hundred loci across all fishes or a thousand loci from acanthomorph fishes, no bait set exists to enrich large numbers (>1000 loci) of ultraconserved nuclear loci from ostariophysans, the second largest actinopterygian superorder.
2. In this manuscript, we describe how we designed a bait set to enrich 2,708 ultraconserved nuclear loci from ostariophysan fishes by combining an existing genome assembly with low coverage sequence data collected from two ostariophysan lineages.
3. We perform a series of enrichment experiments using this bait set across the ostariophysan Tree of Life, from the deepest splits among the major groups (>150 MYA) to more recent divergence events that have occured during the last 50 million years.
4. Our results demonstrate that the bait set we designed is useful for addressing phylogenetic questions from the origin of crown ostariophysans to more recent divergence events, and our *in silico* results suggest that this bait set may be useful for addressing evolutionary questions in closely related groups of fishes, like Clupeiformes.

## Introduction

Target enrichment of highly conserved, phylogenetically informative loci (Faircloth *et al*., 2012) has helped reconstruct and study the evolutionary history of organismal groups ranging from cnidaria and arthropods to vertebrate clades such as birds and snakes (Moyle *et al*., 2016; Streicher and Wiens, 2016; Branstetter *et al*., 2017; Quattrini *et al*., 2018). Among fishes, researchers have designed enrichment bait sets that can collect data from hundreds of loci shared among a majority of ray-finned fishes (Actinopterygii; 33,444 species) (Faircloth *et al*., 2013) or more than one thousand loci shared among actinopterygian subclades (Alfaro *et al*., 2018) like the group of spiny-finned fishes that dominates the world’s oceans (Acanthomorpha; 19,244 species). However, no target enrichment bait set exists that is tailored to collect sequence data from conserved loci shared by ostariophysan fishes, which constitute the second largest actinopterygian superorder (Ostariophysi; 10,887 species).

This ostariophysan radiation (Figure 1) has produced the majority (~70%) of the world’s freshwater fishes and includes catfishes, the milkfish, tetras, minnows, electric knifefishes, and their allies. The evolutionary success of ostariophysans may stem from shared derived possession of an alarm substance called Schreckstoff (von Frisch, 1938) and/or a remarkable modification of the anterior vertebral column known as the Weberian apparatus (Weber, 1820; Rosen *et al*., 1970), which enhances hearing by transmitting sound vibrations from the swim bladder to the ear. Morphological (Fink and Fink, 1981; Fink and Fink, 1996) and molecular studies (Nakatani *et al*., 2011; Betancur-R *et al*., 2013; Arcila *et al*., 2017; Chakrabarty *et al*., 2017) have demonstrated monophyly of the clade and provided numerous hypotheses of relationships among the five ostariophysan orders (reviewed in Chakrabarty *et al*., 2017; Arcila *et al*., 2017). Because several of these hypotheses disagree substantially, major questions about ostariophysan evolution remain unresolved. For example, some studies suggest that Siluriformes (catfishes) and Gymnotiformes (electric knifefishes) are not each other’s closest relatives, which would imply that the electroreceptive capacities of these two orders evolved independently. Other studies have suggested non-monophyly of Characiformes (Chakrabarty *et al*., 2017), which implies a more complicated pattern of evolution in the morphology and development of oral dentition and anatomical systems in this group, as well as suggesting an alternative biogeographical hypothesis to the classical Gondwanan vicariance model (Lundberg, 1993; Sanmartín and Ronquist, 2004). A similar debate concerns the composition of the immediate outgroups to Ostariophysi (see discussion in Lavoué *et al*., 2014), which involve the enigmatic marine family Alepocephalidae (slickheads), as well as the world’s diverse radiation of Clupeiformes (herrings and anchovies), a taxonomic order long allied to Ostariophysi on the basis of anatomical and molecular evidence (Lecointre, 1995).

**Figure 1.**
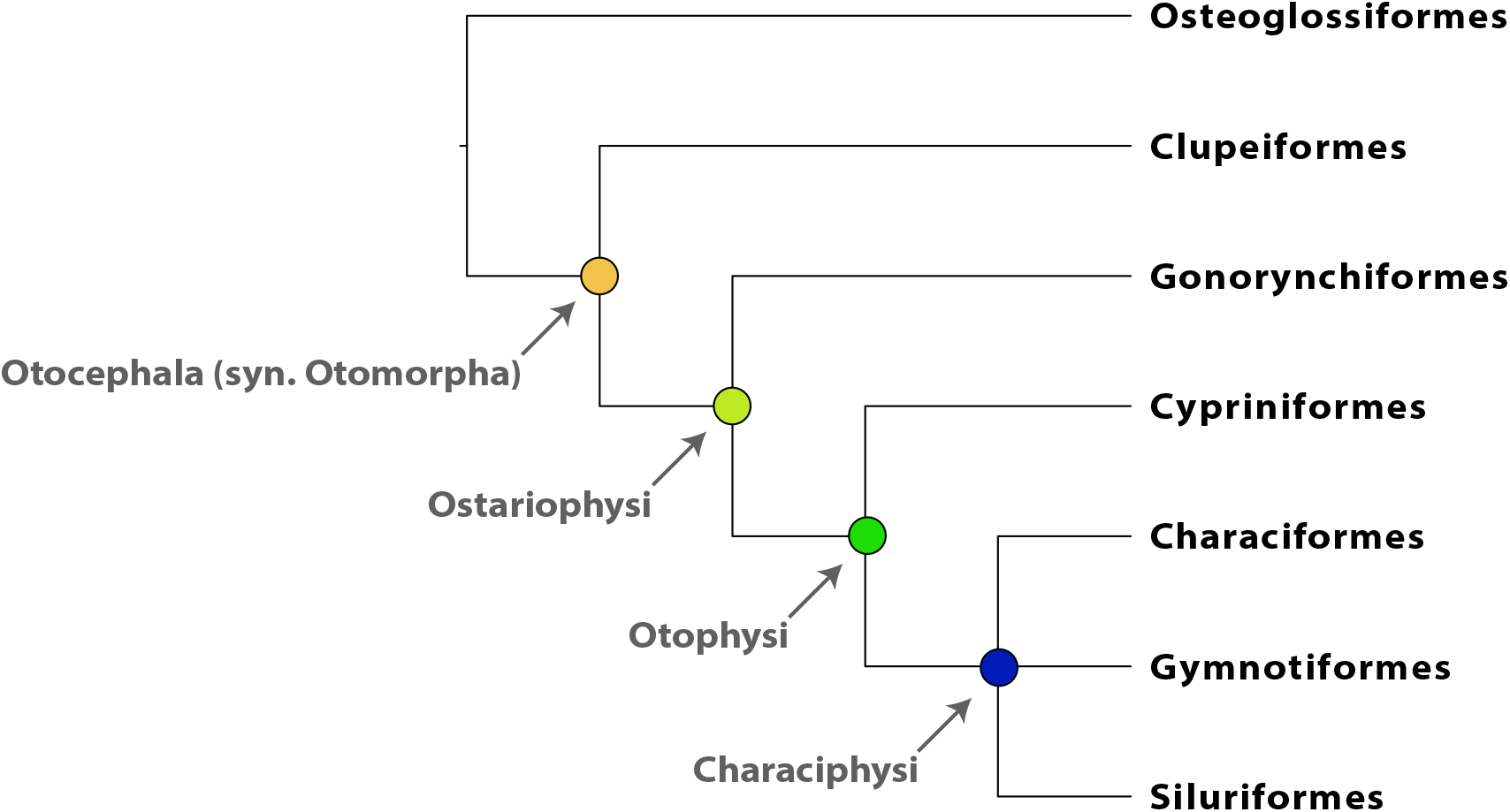
A depiction of relationships among the groups comprising Otocephala and the taxonomic nomenclature used to refer to otocephalan subclades.

Though molecular and morphological hypotheses of interfamilial and intergeneric relationships have been advanced within each of the five ostariophysan orders, substantial work remains before our understanding of the evolutionary history of ostariophysans will rival that of the best studied acanthomorph groups, such as cichlids (Brawand *et al*., 2014; Malinsky *et al*., 2018). The majority of previous work among ostariophysans has involved parsimony analysis of osteological characters or model-based analysis of multilocus Sanger datasets, with even the largest molecular studies (e.g. Schönhuth *et al*., 2018) including less than 15% of the species diversity in the targeted clades. At the genome scale, ostariophysans have been included in studies sampling across the diversity of ray-finned fishes (e.g. Faircloth *et al*., 2013; Hughes *et al*., 2018), while studies focusing on Ostariophysi have only recently begun to appear (Arcila *et al*., 2017; Chakrabarty *et al*., 2017; Dai *et al*., 2018). However, these genome-scale projects have sampled less than 1% of total ostariophysan species diversity and have only begun to address questions of interrelationships among families or genera. A robust and well-documented approach to collect a large number of nuclear loci across ostariophysan orders and appropriate outgroups will accelerate our ability to conduct taxon-rich studies of phylogenetic relationships within and across the group and allow us to synthesize these data into a more complete and modern picture of ostariophysan evolution than previously possible.

Here, we describe the design of an enrichment bait set that targets 2,708 conserved, nuclear loci shared among ostariophysan fishes, and we empirically demonstrate how sequence data collected using this bait set can resolve phylogenetic relationships at several levels of divergence across the ostariophysan Tree of Life, from the deepest splits among ostariophysan orders and their outgroup(s) (Otocephala, crown age 210-178 MYA; (Hughes *et al*., 2018)) to more recent divergence events among lineages comprising the Gymnotiformes (crown age 86-43 MYA) or Anostomoidea (crown age within 76-51 MYA; (Hughes *et al*., 2018)). An earlier manuscript (Arcila *et al*., 2017) developed a bait set targeting 1,068 exon loci shared among otophysans, one of the ostariophysan subclades that includes Characiformes, Cypriniformes, Gymnotiformes, and Siluriformes (Figure 1). The bait set that we describe differs from that of Arcila *et al.* (2017) by targeting a larger number of loci that includes coding and non-coding regions shared among a larger and earlier diverging clade (i.e., ostariophysans and their proximate outroup(s)). As with most bait sets targeting conserved loci shared among related groups, the designs are generally complementary rather than incompatible, and researchers can easily combine loci targeted by both designs to accomplish their research objectives.

## MATERIALS AND METHODS

### Conserved element identification and bait design strategy

When we began this project, few genome assemblies other than zebrafish (*Danio rerio*) were available to represent the diversity of ostariophysans. To identify conserved regions shared among species within this group and design a target enrichment bait set to collect data from these regions, we followed the general steps outlined in (Faircloth, 2017). This means that we needed to identify a “base” genome (*D. rerio*; NCBI Genome GCA_000002035.2) against which to align low coverage sequencing data from exemplar species representing the diversity of ostariophysan lineages. We also needed to identify several exemplar lineages from which to collect low coverage genome sequencing data. Our exemplar taxon selection strategy was to identify and purchase commercially available fish species from orders related to Cypriniformes (to which *D. rerio* belongs) because they were easy to “collect” and provided a ready source of high quality DNA. As a result, we selected *Apteronotus albifrons* and *Corydoras paleatus* as our exemplar taxa. These taxa represent the orders Gymnotiformes (electric knifefishes) and Siluriformes (catfishes), respectively, and together with *D. rerio*, span three of the five orders within Ostariophysi.

### Low coverage genome sequencing

We purchased specimens of *A. albifrons* and *C. paleatus* from commercial wholesalers in the Los Angeles, CA area, collected tissues following protocols approved by the University of California Los Angeles Institutional Animal Care and Use Committee (Approval 2008-176-21), and extracted DNA from each tissue using a commercial kit following the manufacturer’s instructions (DNeasy, Qiagen N.V.). After extraction, we quantified 2 μL of DNA using a Qubit fluorometer (Invitrogen, Inc.) following the manufacturer’s protocol, and we visualized 50-100 ng of each extract by electrophoresis through 1.5% (w/v) agarose gel in TBE or TAE. Following this quality check, we prepared 100 μL (~10 ng/μL) aliquots of extracted DNA, and we sheared each sample to 300–600 bp in length using 5–10 cycles of sonication (High; 30 s on; 90 s off) on a BioRuptor (Diagenode, Inc.). We prepared single-indexed sequencing libraries from 0.5–1.0 μg sheared DNA extracts using a commercial library preparation kit (Kapa Biosystems, Inc.) and a set of custom-indexed sequencing adapters (Faircloth and Glenn, 2012). Following library preparation, we size-selected the *C. paleatus* library to span a range of 200–300 bp using agarose-gel-based size selection. We did not size-select the *A. albifrons* library. We amplified both libraries using 6–10 cycles of PCR, and we purified library amplifications using SPRI beads (Rohland and Reich, 2012). Following purification, we checked the insert size distribution of each library using a BioAnalyzer (Agilent, Inc.), and we quantified libraries using a commercial qPCR quantification kit (Kapa Biosystems, Inc.). We ran each library on a separate lane of Illumina, paired-end, 100 bp sequencing (PE100) by combining each library into a pool of unrelated (and differently indexed) samples, and we sequenced each library pool using an Illumina HiSeq 2500 at the UCLA Neuroscience Genomics Core (UNGC). We demultiplexed the sequencing data using *bcl2fastq* 1.8.4 and allowed one base pair mismatches between the expected and observed indexes (the index sequences we used were robust to ≤ 3 insertion, deletion, or substitution errors).

### Exemplar species validation

We validated the species identification of each sample by aligning FASTQ reads to a related mtDNA genome using *bwa mem* v0.7.17 (Li, 2013), reducing the resulting BAM file to aligning reads using *samtools* v0.1.18 (Li *et al*., 2009), and converting the BAM file of aligned reads back to paired FASTQ reads using *bedtools v2.17.0* (Quinlan and Hall, 2010). We assembled the resulting FASTQ data using *spades v3.10.1* (Nurk *et al*., 2013) with read correction, a kmer length of 55, and the ‘--careful’ assembly option. From the contig that resulted (which was either equal to or slightly shorter than the general mtDNA sequence length for vertebrates), we used a program within the *phyluce* package (Faircloth, 2015) to extract the portions of each contig that were similar to COI sequences from *Apteronotus* (NCBI GenBank AB054132.1:5453-7012) and *Corydoras* (NCBI GenBank JN988809.1). We then matched these extracted COI sequences against the Species Level Barcode Records in the BOLD Systems Database (http://www.boldsystems.org; search performed August 2018). For *A. albifrons*, the top hit (100% sequence identity) was *A. albifrons* (NCBI GenBank AB054132.1; (Saitoh *et al*., 2003)), and for *C. paleatus*, the top publicly available hit (99.85% sequence identity) was *C. paleatus* (NCBI GenBank JX111734.1; (Rosso *et al*., 2012)).

### Conserved element identification and bait design

To identify conserved elements shared among the genomes of *D. rerio, A. albifrons*, and *C. paleatus*, we followed the general workflow described in Faircloth (2017). Specifically, we downloaded the genome assembly of *D. rerio* (hereafter danRer7; NCBI GCA_000002035.2) and aligned the low-coverage, raw reads generated from *A. albifrons* and *C. paleatus* (hereafter “exemplar taxa”) to this reference genome using *stampy* v1.0.21 with the substitution rate set to 0.05. After alignment, we used *samtools* to convert SAM files to BAM format, sort the BAM files, and reduce sorted BAM files to only those reads mapping from the exemplar taxa to the base genome. Using *bedtools*, we converted BAM files to BED coordinates, merged overlapping intervals into a single interval, and computed the intersection of merged intervals among all three taxa. Then, we used programs available as part of the *phyluce* package to filter sequences < 100 bp from the BED files; we extracted the shared, overlapping regions ≥ 100 bp from the danRer7 genome assembly; and we filtered any extracted regions including > 25% masked bases. We then used *phyluce* and *lastz* (Harris, 2007) to search for duplicate hits among these extracted regions, identifying as duplicate two regions that aligned to one another over > 25% of their length with > 85% sequence identity. We used *phyluce* to filter these duplicate hits from the set. These filtered regions represented reasonably long (≥ 100 bp), conserved loci shared among all three taxa. We then ran another filtering step to remove loci from this set that were < 150 bp in length and < 10,000 bp from one another as indicated by their alignment position in the danRer7 genome assembly. We ran a secondary screen for potential duplication against this reduced list using the same parameters as before. Then, we used a program from *phyluce* to design enrichment baits targeting these loci. We attempted to select at least two bait sequences per locus where baits overlapped by 80 bp, and we removed baits from the resulting set with GC content outside the range of 30% to 70%. We used *phyluce* and *lastz* to search for and remove any other baits from this candidate set that were > 50% identical over > 50% of their length. This produced the final ostariophysan bait design file (“ostariophysan bait set” hereafter) which we submitted to Arbor Biosciences, Inc. in order to synthesize myBaits custom target enrichment kits which we used in the empirical tests described below.

### Empirical sequence data collection overview

To test the utility of the resulting bait set for ostariophysan phylogenetics, we designed several experiments that spanned the breadth of species diversity (Table 1) and divergence times in this group. Different research groups performed target captures spanning a range of subclade ages from young (<50 MYA) to old (~200 MYA): Gymnotiformes (crown age 83-46 MYA (Hughes *et al*., 2018)), Anostomoidea (a characiform subclade that includes headstanders and detritivorous characiforms; crown age falls within 76-51 MYA (Hughes *et al*., 2018)), Loricarioidei (armored catfishes; crown age 125 MYA (Rivera-Rivera and Montoya-Burgos, 2017)), and the Characiformes *sensu lato* (tetras and allies; crown age 133-112 MYA (Hughes *et al*., 2018). We then combined data from several species within each group with additional enrichments from outgroup lineages and conserved loci harvested from available genome sequences to create a data set spanning Otocephala, a diverse teleostean clade that includes ostariophysans and clupeomorphs (sardines, herrings and allies; crown age 210-178 MYA).

**Table 1.**
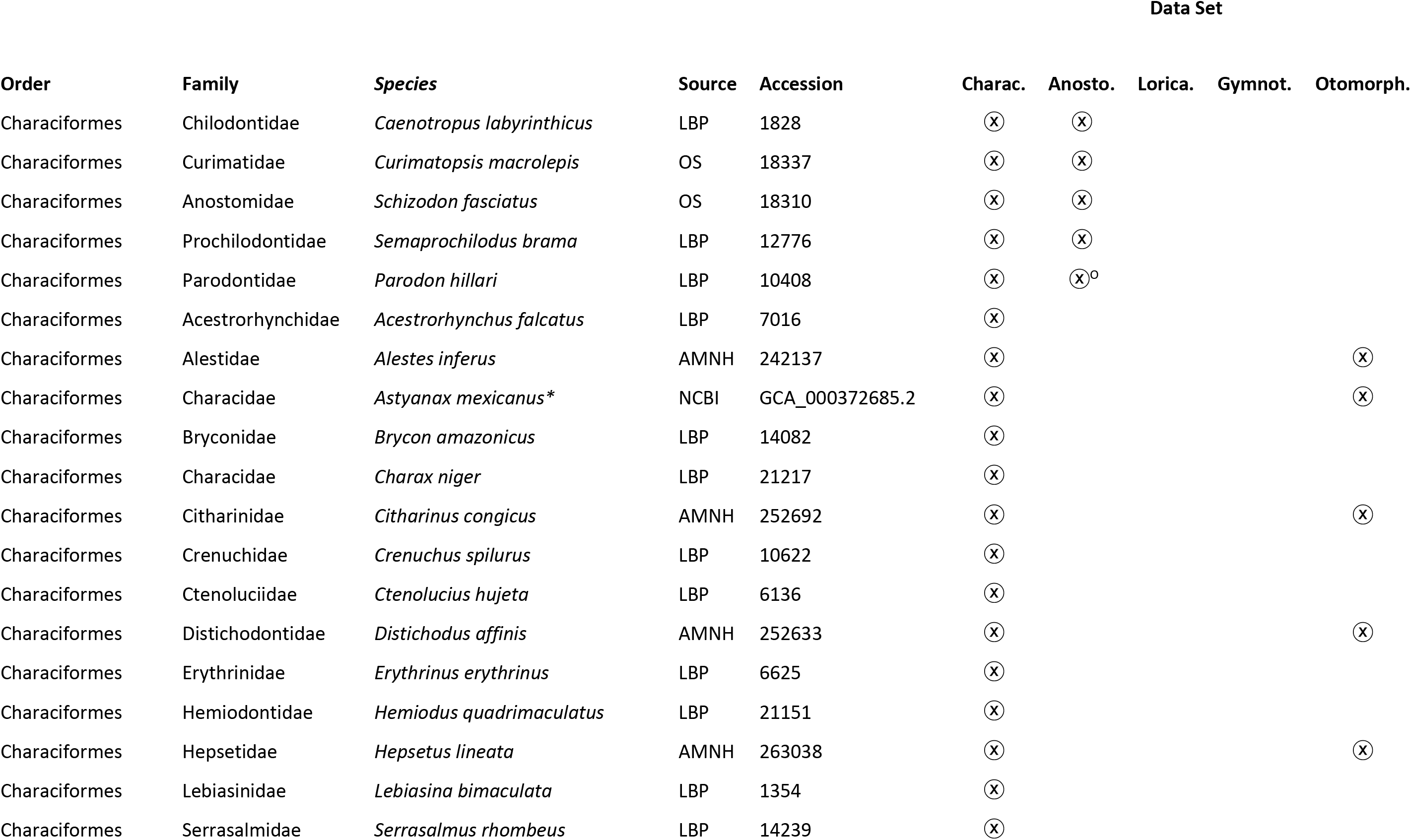
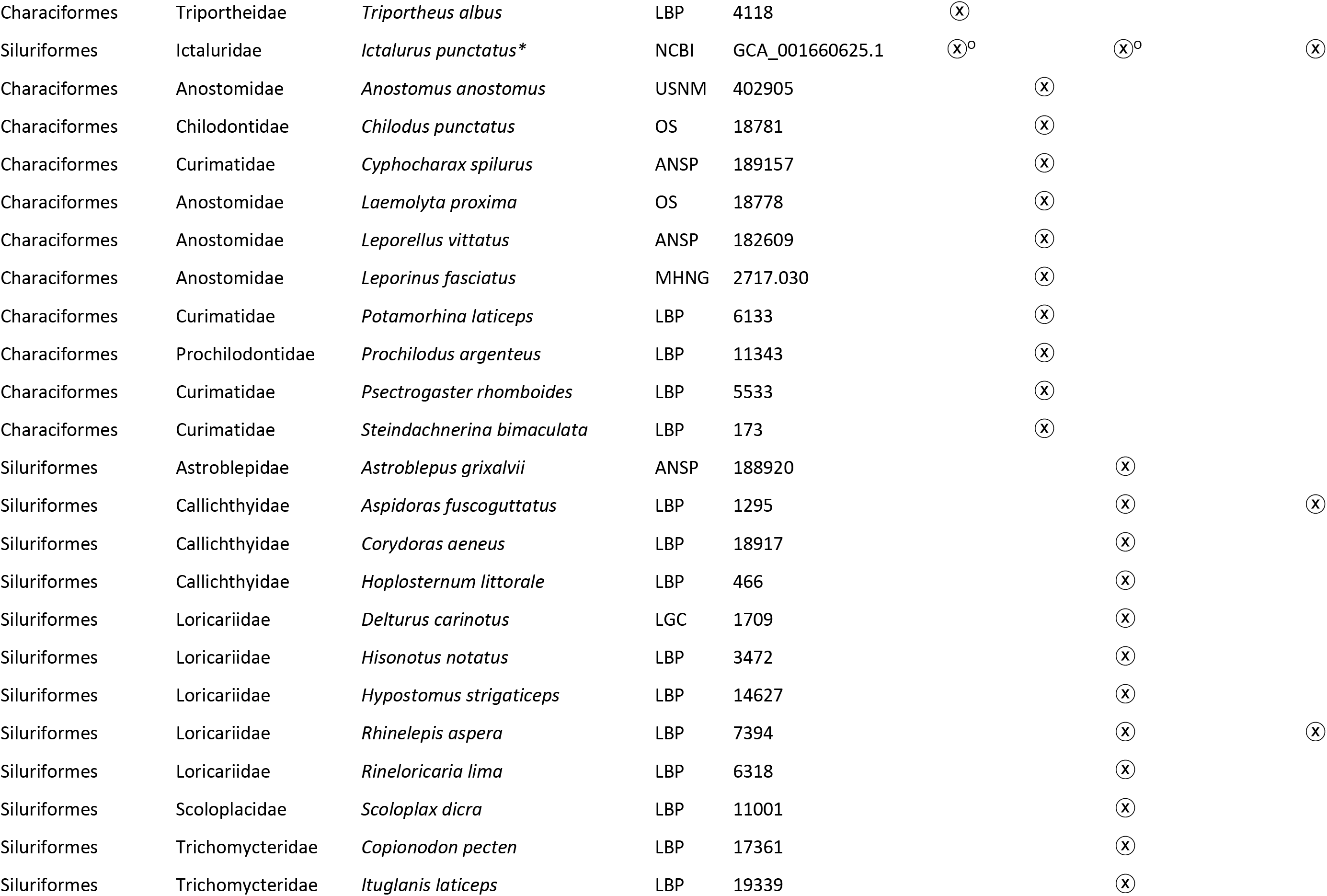
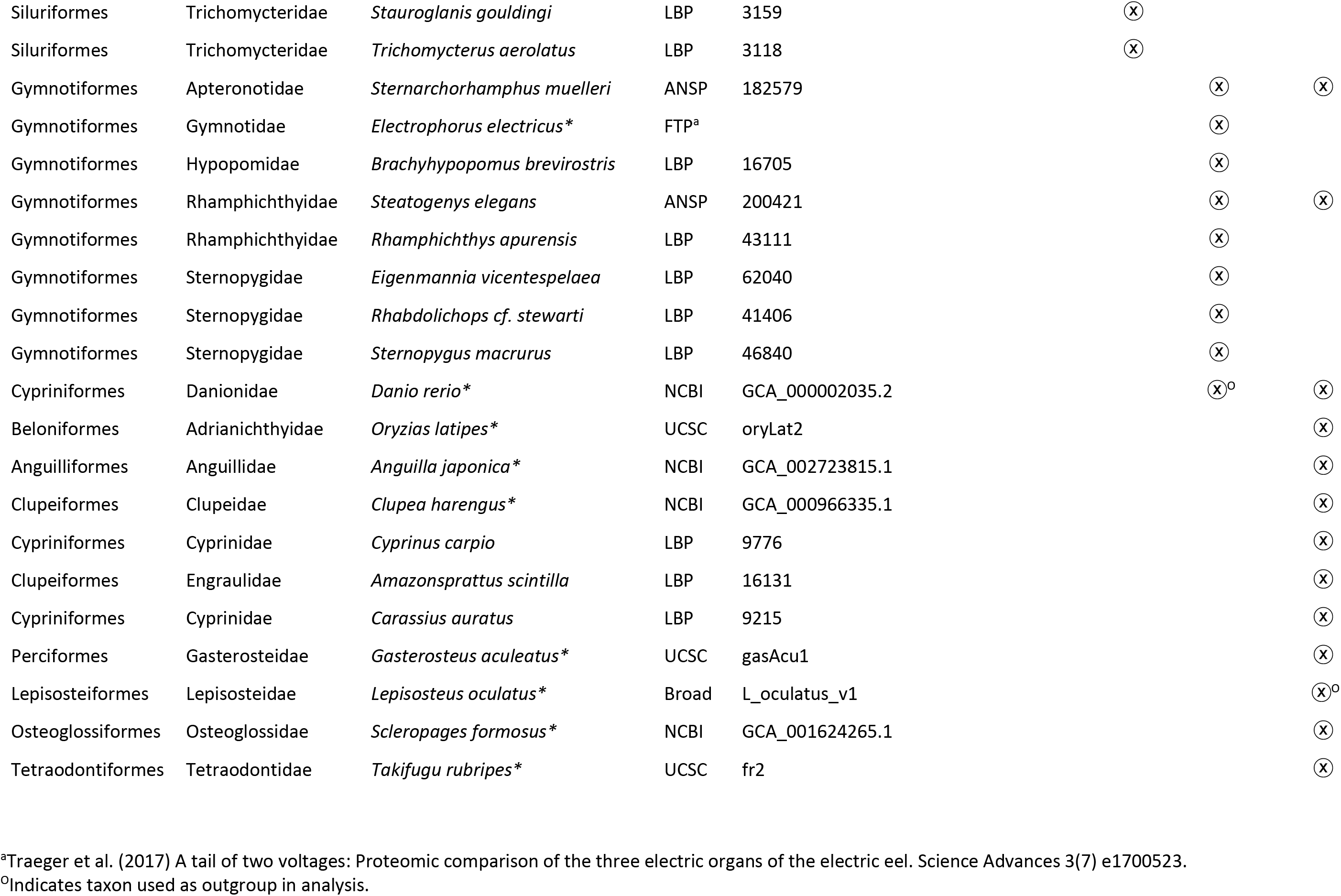
Order, family, species, source, accession and data set membership of fish species from which we enriched or bioinformatically harvested UCE loci. Charac. = Characiformes, Anosto. = Anostomoidea, Lorica. = Loricarioidei, Gymnot. = Gymnotiformes, Otomorph. = Otomorpha.

#### Library preparation and enrichment from Gymnotiformes

We extracted DNA from vouchered museum specimens (Table 1) for taxa in the Gymnotiformes data set using DNeasy Tissue kits (Qiagen N.V.). We quantified DNA using a Qubit fluorometer (Invitrogen, Inc.), checked the quality of extracted DNA by visualization using gel electrophoresis (1.5% agarose w/v), and sheared 300–500 ng of DNA extracts to a target size of 400–600 bp using 20 cycles of sonication (15 s on; 30 s off) on an EpiSonic Multi-Functional Bioprocessor (Epigentek Group, Inc). We prepared sequencing libraries using Kapa Hyper Prep Kits (Kapa Biosystems, Inc). We incorporated sample-specific dual-indexes to library fragments by ligating universal iTru stubs to DNA and performing 10 cycles of PCR with iTru dual-indexed primers (Glenn *et al*., 2016) and Kapa HiFi Hotstart ReadyMix polymerase. We cleaned indexed libraries using SPRI beads (Rohland and Reich, 2012), quantified clean libraries using a Qubit fluorometer, and created pools of 8 libraries at 62.5 ng each (500 ng total). We diluted the ostariophysan bait set synthesized by Arbor Biosciences at a 1:6 ratio, and we enriched library pools following the MYBaits kit v3.0 protocol. We PCR-recovered enriched library pools using 16 PCR cycles, Kapa HiFi HotStart Taq, and the Illumina P5/P7 primer pair. We purified the resulting product using SPRI beads; quantified enriched, clean libraries using a Qubit fluorometer; and combined libraries at equimolar concentrations with other enriched pools having different indexes. We then sequenced enriched libraries by combining them with other library pools having different indexes in two separate runs of a PE150 Illumina NextSeq 300 (University of Georgia Genomics Facility) or a PE150 Illumina HiSeq 3000 (Oklahoma Medical Research Foundation). We received demultiplexed sequence data from the sequencing facilities.

#### Library preparation and enrichment from Anostomoidea

We extracted DNA from vouchered museum samples (Table 1) for the taxa in the Anostomoidea data set using either a DNeasy Tissue kit or a modified NaCl extraction protocol adapted from Lopera-Barrero *et al.* (2008). Following extraction, we quantified DNA using a Qubit fluorometer and visualized 50–100 ng of each extract by electrophoresis through 1.5% (w/v) agarose gel in TBE. We prepared 100 μL aliquots (~10 ng/μL) of each sample for sonication, and we sheared samples to 300–600 bp using 5–10 cycles of sonication (High; 30 s on; 90 s off) on a BioRuptor (Diagenode, Inc.). We prepared sequencing libraries from sheared DNA using a commercial library preparation kit (Kapa Biosystems, Inc.). We indexed individual libraries using either a set of single, custom indexes (Faircloth and Glenn, 2012) or the iTru dual-indexing approach (Glenn *et al*., 2016). We PCR-amplified libraries using Kapa HiFi HotStart ReadyMix polymerase, 14 cycles of PCR, and the manufacturer’s recommended thermal profile. We purified library amplifications using SPRI beads, quantified libraries with a Qubit fluorometer, and concentrated libraries to 147 ng/μL with a Speed-Vac prior to preparing pools of 8 libraries at 62.5 ng each (500 ng total). We enriched libraries by diluting the ostariophysan bait set (1:5) and following the procedure described for Gymnotiformes. We combined enriched library pools with other libraries having different indexes, and we sequenced the pool of pooled libraries using PE125 sequencing on an Illumina HiSeq 2000 in the Center for Genome Research and Biocomputing at Oregon State University.

#### Library preparation and enrichment from Loricaroidei and Characiformes

We extracted DNA from vouchered museum samples (Table 1) for the taxa in the Loricaroidei and Characiformes data sets using DNeasy Tissue kits. Subsequent library preparation and enrichment steps were performed by Arbor Biosciences, Inc. (AB). AB staff quantified and sheared DNA extracts with a Q800R (QSonica, Inc.) instrument and performed dual SPRI size selection to produce sheared DNAs having a modal length of approximately 500 bp. AB staff then prepared sequencing libraries using NEBNext Ultra DNA Library Preparation Kits (New England Biolabs, Inc.). Following ligation, AB staff amplified libraries using dual P7 and P5 indexing primers (Kircher *et al*., 2012), KAPA HiFi HotStart ReadyMix (Kapa Biosystems, Inc.), six PCR cycles, and the manufacturer’s recommended thermal profile. AB staff purified libraries with SPRI beads, quantified libraries with the Quant-iT Picogreen dsDNA Assay kit (ThermoFisher, Inc.), prepared pools of 8 libraries at 100 ng each (800ng total input material), and enriched libraries using an undiluted ostariophysan bait set following following the MYBaits kit v3.0 protocol. After capture, AB staff resuspended enriched, bead-bound library pools in the recommended solution and PCR-recovered enriched library pools using 10 PCR cycles, Kapa HiFi polymerase, and the Illumina P5/P7 primer pair. AB staff purified PCR-recovered libraries using SPRI beads, quantified enriched, clean libraries with PicoGreen, and combined enriched pools with other libraries having different indexes at the desired ratio prior to PE100 sequencing using an Illumina HiSeq 2500.

### Sequence data quality control and assembly

After sequencing, we received FASTQ data from each sequencing provider, and we removed adapters and trimmed the sequence data for low quality bases using *illumiprocessor* (https://illumiprocessor.readthedocs.io/) which is a wrapper around Trimmomatic (Bolger *et al*., 2014). We assembled trimmed reads using a *phyluce* wrapper around the Trinity assembly program (Grabherr *et al*., 2011). Before creating data sets for phylogenetic processing, we integrated the sequence data collected *in vitro* with those collected *in silico.*

### In-silico sequence data collection

We used computational approaches to extract data from 11 fish genome assemblies available from UCSC, NCBI, and other sites (Table 1). We identified and extracted UCE loci that matched the ostariophysan bait set using *phyluce* and a standardized workflow (https://phyluce.readthedocs.io/en/latest/tutorial-three.html), except that we adjusted the sequence coverage value to 67% and the sequence identity parameter to 80%. After locus identification, we sliced UCE loci ± 500 bp from each genome and output those slices into FASTA files identical to the FASTA files generated from assemblies of the samples we processed *in vitro.* After harvesting the *in silico* data, we merged these with the *in vitro* data and processed both simultaneously.

### UCE identification, alignment, and phylogenetic analyses

We used a standard workflow (https://phyluce.readthedocs.io/en/latest/tutorial-one.html) and programs within *phyluce* to identify and filter non-duplicate contigs representing conserved loci enriched by the ostariophysan bait set (hereafter UCEs). Then, we used lists of taxa to create one data set for each taxonomic group outlined in Table 1, and we extracted FASTA data from the UCE contigs enriched for group members. We exploded these data files by taxon to compute summary metrics for UCE contigs, and we used *phyluce* to generate mafft v.7 (Katoh and Standley, 2013) alignments of all loci. We trimmed alignments using *trimAL* (Capella-Gutierrez *et al*., 2009) and the ‘-automated1’ routine, and we computed alignment statistics using *phyluce.* We then generated 75% complete data matrices for all data sets, and we computed summary statistics across each 75% complete matrix. We concatenated alignments using *phyluce*, and we conducted maximum likelihood (ML) tree and bootstrap replicate searches with the GTRGAMMA site rate substitution model using RAxML (v8.0.19). We used the ‘-autoMRE’ function of RAxML to automatically determine the bootstrap replicate stopping point. Following best and bootstrap ML tree searches, we added bootstrap support values to each tree using RAxML. We did not run Bayesian or coalescent-based analyses because we were interested in determining whether this bait set produced reasonable results at the levels of divergence examined rather than exhaustively analyzing the evolutionary relationships among the taxa included.

## RESULTS

We collected an average of 3.47 M reads from enriched libraries (Table 2), and we assembled these reads into an average of 18,048 contigs having a mean length of 440 bp (Table 3). After searching for enriched, conserved loci among the contig assemblies, we identified an average of 1,446 targeted, conserved loci per library (range 525–1882; Table 4) having a mean length of 666 bp per locus. From these loci, we created five different data sets (Table 1) that spanned the diversity of relationships within ostariophysans and extended beyond this clade to include Clupeiformes and other distantly related lineages (the Otocephala data set). We describe specific results from each of these data sets below.

**Table 2.** The number of reads sequenced and summary statistics of trimmed reads for each DNA library from which we enriched UCEs using the ostariophysan bait set.

| Order | Family | Species | Reads | Mean<br>Trimmed<br>Length | 95 CI<br>Trimmed<br>Length | Min | Max | Median |
| --- | --- | --- | --- | --- | --- | --- | --- | --- |
| Characiformes | Acestrorhynchidae | *Acestrorhynchus falcatus* | 2,046,124 | 98.24 | 0.01 | 40 | 100 | 100 |
| Characiformes | Alestidae | *Alestes inferus* | 1,847,106 | 118.76 | 0.01 | 40 | 125 | 125 |
| Characiformes | Anostomidae | *Anostomus anostomus* | 1,357,565 | 95.15 | 0.01 | 40 | 101 | 101 |
| Characiformes | Anostomidae | *Laemolyta proxima* | 3,330,657 | 96.09 | 0.01 | 40 | 101 | 101 |
| Characiformes | Anostomidae | *Leporellus vittatus* | 3,734,248 | 96.08 | 0.01 | 40 | 101 | 101 |
| Characiformes | Anostomidae | *Leporinus fasciatus* | 1,054,553 | 95.53 | 0.01 | 40 | 101 | 101 |
| Characiformes | Anostomidae | *Schizodon fasciatus* | 6,350,444 | 96.72 | 0.00 | 40 | 101 | 101 |
| Characiformes | Bryconidae | *Brycon amazonicus* | 3,916,413 | 120.13 | 0.01 | 40 | 125 | 125 |
| Characiformes | Characidae | *Charax niger* | 4,557,912 | 118.99 | 0.01 | 40 | 125 | 125 |
| Characiformes | Chilodontidae | *Caenotropus labyrinthicus* | 8,097,664 | 98.53 | 0.00 | 40 | 101 | 101 |
| Characiformes | Chilodontidae | *Chilodus punctatus* | 5,695,056 | 95.90 | 0.01 | 40 | 101 | 101 |
| Characiformes | Citharinidae | *Citharinus congicus* | 5,077,942 | 119.74 | 0.01 | 40 | 125 | 125 |
| Characiformes | Crenuchidae | *Crenuchus spilurus* | 2,693,174 | 94.68 | 0.01 | 40 | 101 | 101 |
| Characiformes | Ctenoluciidae | *Ctenolucius hujeta* | 3,758,899 | 119.84 | 0.01 | 40 | 125 | 125 |
| Characiformes | Curimatidae | *Curimatopsis macrolepis* | 5,082,531 | 96.24 | 0.01 | 40 | 101 | 101 |
| Characiformes | Curimatidae | *Cyphocharax spilurus* | 6,010,208 | 95.92 | 0.01 | 40 | 101 | 101 |
| Characiformes | Curimatidae | *Potamorhina laticeps* | 1,559,277 | 95.61 | 0.01 | 40 | 101 | 101 |
| Characiformes | Curimatidae | *Psectrogaster rhomboides* | 10,674,224 | 98.61 | 0.00 | 40 | 101 | 101 |
| Characiformes | Curimatidae | *Steindachnerina bimaculata* | 1,656,298 | 93.52 | 0.01 | 40 | 101 | 101 |
| Characiformes | Distichodontidae | *Distichodus affinis* | 5,431,103 | 119.44 | 0.01 | 40 | 125 | 125 |
| Characiformes | Erythrinidae | *Erythrinus erythrinus* | 3,639,480 | 118.76 | 0.01 | 40 | 125 | 125 |
| Characiformes | Hemiodontidae | *Hemiodus quadrimaculatus* | 1,631,554 | 117.11 | 0.01 | 40 | 125 | 125 |
| Characiformes | Hepsetidae | *Hepsetus lineata* | 4,790,584 | 119.77 | 0.01 | 40 | 125 | 125 |
| Characiformes | Lebiasinidae | *Lebiasina bimaculata* | 3,046,673 | 118.68 | 0.01 | 40 | 125 | 125 |
| Characiformes | Parodontidae | *Parodon hillari* | 4,561,066 | 98.14 | 0.00 | 40 | 101 | 101 |
| Characiformes | Prochilodontidae | *Prochilodus argenteus* | 1,092,497 | 95.65 | 0.01 | 40 | 101 | 101 |
| Characiformes | Prochilodontidae | *Semaprochilodus brama* | 8,465,340 | 98.87 | 0.00 | 40 | 101 | 101 |
| Characiformes | Serrasalminae | *Serrasalmus rhombeus* | 1,662,084 | 118.92 | 0.01 | 40 | 125 | 125 |
| Characiformes | Triportheidae | *Triportheus albus* | 1,131,229 | 118.80 | 0.02 | 40 | 125 | 125 |
| Clupeiformes | Engraulidae | *Amazonsprattus scintilla* | 1,666,800 | 115.99 | 0.01 | 40 | 125 | 125 |
| Cypriniformes | Cyprinidae | *Cyprinus carpio* | 1,232,988 | 120.37 | 0.01 | 40 | 125 | 125 |
| Cypriniformes | Cyprinidae | *Carassius auratus* | 2,828,778 | 119.19 | 0.01 | 40 | 125 | 125 |
| Gymnotiformes | Apteronotidae | *Sternarchorhamphus muelleri* | 9,914,493 | 129.60 | 0.01 | 40 | 151 | 149 |
| Gymnotiformes | Hypopomidae | *Brachyhypopomus brevirostris* | 2,493,236 | 127.80 | 0.02 | 40 | 151 | 149 |
| Gymnotiformes | Rhamphichthyidae | *Rhamphichthys apurensis* | 3,358,202 | 124.09 | 0.02 | 40 | 151 | 149 |
| Gymnotiformes | Rhamphichthyidae | *Steatogenys elegans* | 4,932,755 | 117.97 | 0.02 | 40 | 151 | 136 |
| Gymnotiformes | Sternopygidae | *Eigenmannia vicentespela* | 4,754,970 | 126.41 | 0.02 | 40 | 151 | 149 |
| Gymnotiformes | Sternopygidae | *Rhabdolichops cf. stewarti* | 5,764,578 | 123.57 | 0.02 | 40 | 151 | 149 |
| Gymnotiformes | Sternopygidae | *Sternopygus macrurus* | 3,667,938 | 128.86 | 0.02 | 40 | 151 | 150 |
| Siluriformes | Astroblepidae | *Astroblepus grimaldi* | 1,227,559 | 118.33 | 0.01 | 40 | 125 | 125 |
| Siluriformes | Callichthyidae | *Aspidoras fuscoguttatus* | 2,309,437 | 98.01 | 0.01 | 40 | 100 | 100 |
| Siluriformes | Callichthyidae | *Corydoras aeneus* | 3,070,805 | 119.78 | 0.01 | 40 | 125 | 125 |
| Siluriformes | Callichthyidae | *Hoplosternum littorale* | 2,602,840 | 98.38 | 0.00 | 40 | 100 | 100 |
| Siluriformes | Loricariidae | *Delturus carinatus* | 823,134 | 120.64 | 0.01 | 40 | 125 | 125 |
| Siluriformes | Loricariidae | *Hisonotus notatus* | 1,869,986 | 98.05 | 0.01 | 40 | 100 | 100 |
| Siluriformes | Loricariidae | *Hypostomus strigaticeps* | 524,859 | 120.55 | 0.02 | 40 | 125 | 125 |
| Siluriformes | Loricariidae | *Rhinelepis aspera* | 1,833,881 | 98.57 | 0.00 | 40 | 100 | 100 |
| Siluriformes | Loricariidae | *Rineloricaria lima* | 1,569,938 | 97.59 | 0.01 | 40 | 100 | 100 |
| Siluriformes | Scoloplacidae | *Scoloplax dicra* | 3,626,939 | 97.38 | 0.00 | 40 | 100 | 100 |
| Siluriformes | Trichomycteridae | *Copionodon pecten* | 2,000,677 | 98.30 | 0.01 | 40 | 100 | 100 |
| Siluriformes | Trichomycteridae | *Ituglanis laticeps* | 3,235,129 | 97.94 | 0.00 | 40 | 100 | 100 |
| Siluriformes | Trichomycteridae | *Stauroglanis gouldingi* | 2,053,042 | 98.09 | 0.01 | 40 | 100 | 100 |
| Siluriformes | Trichomycteridae | *Trichomycterus aerolatus* | 2,820,795 | 97.87 | 0.00 | 40 | 100 | 100 |

**Table 3.**
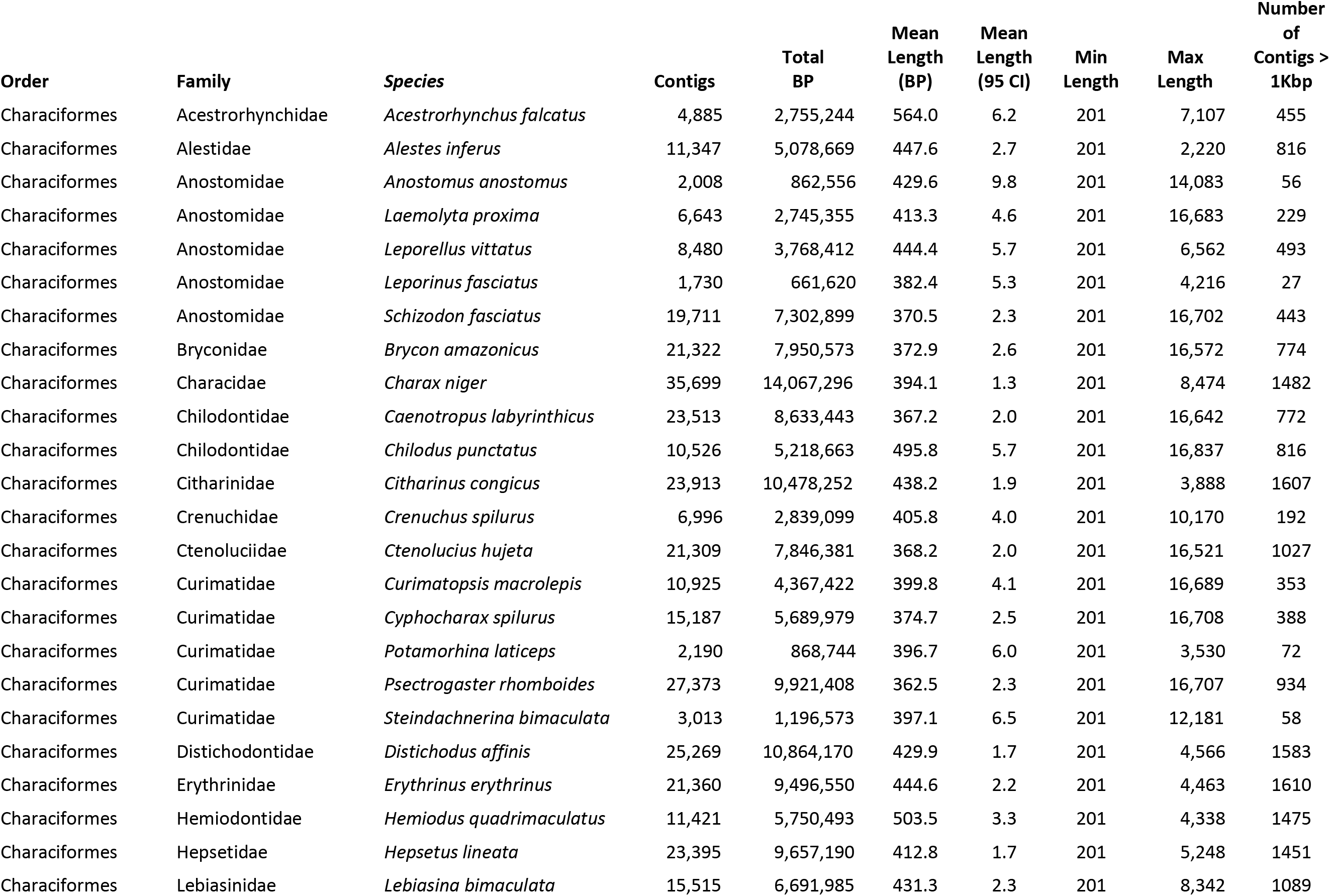
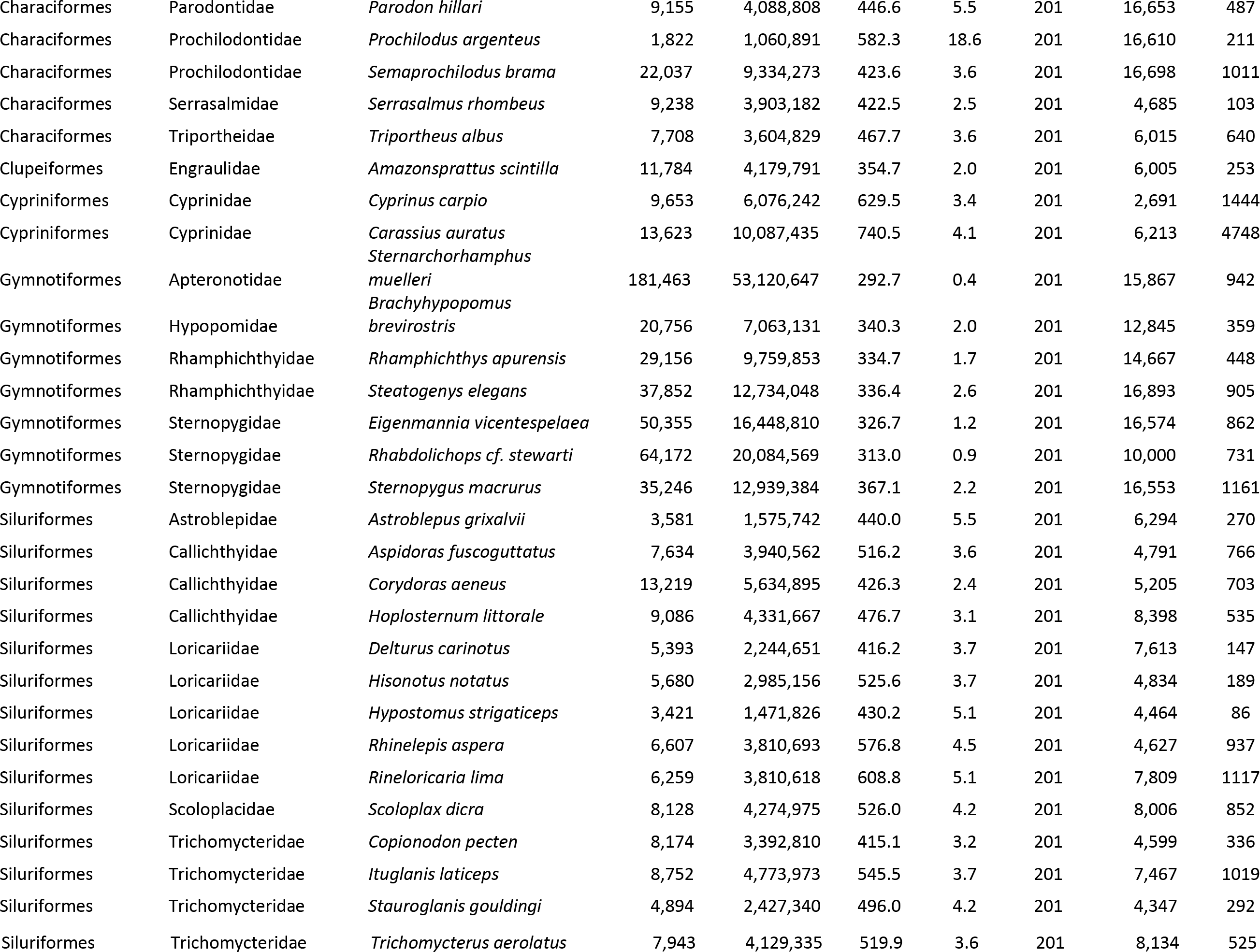
The number of assembled contigs and summary statistics of assembled contigs for each enriched DNA library.

### Gymnotiformes data set

The Gymnotiformes data set (Table 1) was one of two “young” ostariophysan subclades we studied (crown age 83-46 MYA (Hughes *et al*., 2018)). We enriched an average of 1,871 UCE loci from members of this group that averaged 591 bp in length and represented 2,259 of 2,708 loci (83%) that we targeted (Table 4). Alignments generated from these loci contained an average of 7 taxa (range 3–9). After alignment trimming, the 75% matrix contained 1,771 UCE loci including an average of 8 taxa (range 6–9) and having an average trimmed length of 466, a total length of 825,574 characters, and an average of 62 parsimony informative sites per locus. RAxML bootstrap analyses required 50 iterations to reach the MRE stopping point.

**Table 4.** The number of ultraconserved element (UCE) loci identified among the assembled contigs and summary statistics of these UCE loci for all taxa included in this study. Asterisks by species names indicate those taxa for which we included UCE loci harvested in silico.

| Order | Family | Species | UCE<br>Contigs | UCE<br>Contig<br>Mean<br>Length | UCE<br>Contig<br>95 CI | UCE<br>Contig<br>Min<br>Length | UCE<br>Contig<br>Max<br>Length | UCE<br>Contig<br>Median<br>Length | UCE<br>Contigs<br>> 1<br>Kbp |
| --- | --- | --- | --- | --- | --- | --- | --- | --- | --- |
| Anguilliformes | Anguillidae | *Anguilla japonica* * | 897 | 196.1 | 1.8 | 110 | 771 | 187 | 0 |
| Beloniformes | Adrianichthyidae | *Oryzias latipes* * | 969 | 196.9 | 1.7 | 78 | 726 | 190 | 0 |
| Characiformes | Acestrorhynchidae | *Acestrorhynchus falcatus* | 1667 | 713.0 | 6.9 | 201 | 4420 | 728 | 257 |
| Characiformes | Alestidae | *Alestes inferus* | 1788 | 777.5 | 7.1 | 202 | 1551 | 792.5 | 480 |
| Characiformes | Anostomidae | *Anostomus anostomus* | 855 | 432.9 | 16.8 | 201 | 14083 | 387 | 1 |
| Characiformes | Anostomidae | *Laemolyta proxima* | 1280 | 484.6 | 5.1 | 201 | 1075 | 468 | 6 |
| Characiformes | Anostomidae | *Leporellus vittatus* | 1379 | 443.4 | 4.2 | 201 | 1003 | 429 | 1 |
| Characiformes | Anostomidae | *Leporinus fasciatus* | 790 | 386.1 | 5.3 | 201 | 2363 | 371.5 | 1 |
| Characiformes | Anostomidae | *Schizodon fasciatus* | 1613 | 519.8 | 4.7 | 201 | 1173 | 507 | 11 |
| Characiformes | Bryconidae | *Brycon amazonicus* | 1792 | 709.1 | 6.8 | 203 | 5902 | 714 | 251 |
| Characiformes | Characidae | *Astyanax mexicanus* * | 1880 | 1102.6 | 2.9 | 265 | 1647 | 1132.5 | 1626 |
| Characiformes | Characidae | *Charax niger* | 1701 | 851.3 | 8.1 | 201 | 1888 | 878 | 641 |
| Characiformes | Chilodontidae | *Caenotropus labyrinthicus* | 1676 | 672.0 | 11.9 | 201 | 16642 | 637.5 | 249 |
| Characiformes | Chilodontidae | *Chilodus punctatus* | 1579 | 596.5 | 6.3 | 201 | 2453 | 570 | 99 |
| Characiformes | Citharinidae | *Citharinus congicus* | 1721 | 875.2 | 8.0 | 202 | 3765 | 887 | 664 |
| Characiformes | Crenuchidae | *Crenuchus spilurus* | 1195 | 495.0 | 5.2 | 201 | 1323 | 485 | 5 |
| Characiformes | Ctenoluciidae | *Ctenolucius hujeta* | 1594 | 799.4 | 7.0 | 202 | 1423 | 832 | 466 |
| Characiformes | Curimatidae | *Curimatopsis macrolepis* | 1474 | 506.2 | 4.9 | 201 | 1160 | 492 | 7 |
| Characiformes | Curimatidae | *Cyphocharax spilurus* | 1477 | 460.1 | 4.4 | 201 | 1392 | 441 | 4 |
| Characiformes | Curimatidae | *Potamorhina laticeps* | 799 | 357.4 | 4.4 | 201 | 1604 | 335 | 1 |
| Characiformes | Curimatidae | *Psectrogaster rhomboides* | 1465 | 594.1 | 7.8 | 201 | 1734 | 535 | 176 |
| Characiformes | Curimatidae | *Steindachnerina bimaculata* | 868 | 462.1 | 14.8 | 201 | 12181 | 419 | 1 |
| Characiformes | Distichodontidae | *Distichodus affinis* | 1710 | 915.6 | 7.6 | 201 | 2114 | 958.5 | 769 |
| Characiformes | Erythrinidae | *Erythrinus erythrinus* | 1756 | 859.6 | 7.2 | 202 | 2259 | 892 | 673 |
| Characiformes | Hemiodontidae | *Hemiodus quadrimaculatus* | 1836 | 874.8 | 7.6 | 203 | 2066 | 918 | 764 |
| Characiformes | Hepsetidae | *Hepsetus lineata* | 1814 | 918.3 | 8.0 | 202 | 5248 | 955 | 835 |
| Characiformes | Lebiasinidae | *Lebiasina bimaculata* | 1779 | 852.8 | 6.8 | 201 | 1855 | 885 | 629 |
| Characiformes | Parodontidae | *Parodon hillari* | 1472 | 525.1 | 5.5 | 201 | 1098 | 502.5 | 18 |
| Characiformes | Prochilodontidae | *Prochilodus argenteus* | 693 | 365.5 | 23.8 | 201 | 16610 | 320 | 1 |
| Characiformes | Prochilodontidae | *Semaprochilodus brama* | 1671 | 597.8 | 5.7 | 201 | 1408 | 597 | 63 |
| Characiformes | Serrasalminidae | *Serrasalmus rhombeus* | 1830 | 656.2 | 4.4 | 201 | 1283 | 687 | 25 |
| Characiformes | Triportheidae | *Triportheus albus* | 1660 | 753.6 | 6.9 | 201 | 1793 | 774 | 394 |
| Clupeiformes | Clupeidae | *Clupea harengus* * | 1360 | 1118.1 | 3.4 | 188 | 2785 | 1136 | 1246 |
| Clupeiformes | Engraulidae | *Amazonsprattus scintilla* | 867 | 607.3 | 10.1 | 201 | 3038 | 556 | 96 |
| Cypriniformes | Cyprinidae | *Cyprinus carpio* | 780 | 767.3 | 9.7 | 201 | 2599 | 802.5 | 160 |
| Cypriniformes | Danionidae | *Danio rerio* * | 2218 | 227.7 | 1.1 | 141 | 751 | 210 | 0 |
| Cypriniformes | Cyprinidae | *Carassius auratus* | 525 | 960.8 | 17.3 | 202 | 2919 | 1050 | 278 |
| Gymnotiformes | Apteronotidae | *Sternarchorhamphus muelleri* | 1881 | 611.5 | 5.4 | 201 | 5445 | 615 | 54 |
| Gymnotiformes | Gymnotidae | *Electrophorus electricus* * | 1928 | 1152.7 | 1.8 | 388 | 1640 | 1156 | 1877 |
| Gymnotiformes | Hypopomidae | *Brachyhypopomus brevirostris* | 1779 | 531.4 | 8.0 | 201 | 12845 | 528 | 7 |
| Gymnotiformes | Rhamphichthyidae | *Rhamphichthys apurensis* | 1759 | 557.6 | 4.1 | 202 | 1139 | 567 | 9 |
| Gymnotiformes | Rhamphichthyidae | *Steatogenys elegans* | 1771 | 491.6 | 3.8 | 201 | 2194 | 488 | 2 |
| Gymnotiformes | Sternopygidae | *Eigenmannia vicentespela* | 1836 | 582.1 | 4.3 | 202 | 2019 | 593 | 17 |
| Gymnotiformes | Sternopygidae | *Rhabdolichops cf. stewarti* | 1824 | 593.2 | 5.1 | 202 | 4650 | 597 | 30 |
| Gymnotiformes | Sternopygidae | *Sternopygus macrurus* | 1849 | 569.4 | 9.5 | 202 | 16553 | 566 | 9 |
| Lepisosteiformes | Lepisosteidae | *Lepisosteus oculatus* * | 1158 | 202.7 | 1.6 | 125 | 710 | 199 | 0 |
| Osteoglossiformes | Osteoglossidae | *Scleropages formosus* * | 944 | 194.3 | 1.6 | 67 | 690 | 190 | 0 |
| Perciformes | Gasterosteidae | *Gasterosteus aculeatus* * | 1052 | 202.4 | 1.6 | 100 | 650 | 199 | 0 |
| Siluriformes | Astroblepidae | *Astroblepus grimaldi* | 859 | 680.2 | 11.7 | 201 | 1791 | 674 | 186 |
| Siluriformes | Callichthyidae | *Aspidoras fuscoguttatus* | 1525 | 838.1 | 6.9 | 202 | 1630 | 856 | 455 |
| Siluriformes | Callichthyidae | *Corydoras aeneus* | 1411 | 758.1 | 7.3 | 201 | 2179 | 784 | 276 |
| Siluriformes | Callichthyidae | *Hoplosternum littorale* | 1587 | 779.6 | 6.3 | 202 | 1782 | 792 | 324 |
| Siluriformes | Ictaluridae | *Ictalurus punctatus* * | 1862 | 1138.9 | 1.7 | 465 | 1599 | 1144 | 1804 |
| Siluriformes | Loricariidae | *Delturus carinotus* | 1329 | 639.6 | 6.8 | 202 | 1618 | 660 | 86 |
| Siluriformes | Loricariidae | *Hisonotus notatus* | 1491 | 705.6 | 5.6 | 201 | 1556 | 740 | 71 |
| Siluriformes | Loricariidae | *Hypostomus strigaticeps* | 1070 | 538.6 | 6.5 | 201 | 1293 | 532 | 13 |
| Siluriformes | Loricariidae | *Rhinelepis aspera* | 1542 | 855.6 | 7.5 | 202 | 1723 | 879 | 533 |
| Siluriformes | Loricariidae | *Rineloricaria lima* | 1416 | 899.4 | 8.4 | 201 | 1771 | 917 | 579 |
| Siluriformes | Scoloplacidae | *Scoloplax dicra* | 1087 | 821.4 | 9.4 | 203 | 2020 | 825 | 315 |
| Siluriformes | Trichomycteridae | *Copionodon pecten* | 1380 | 742.0 | 6.7 | 201 | 2535 | 764 | 186 |
| Siluriformes | Trichomycteridae | *Ituglanis laticeps* | 1511 | 848.5 | 7.5 | 202 | 1996 | 856 | 491 |
| Siluriformes | Trichomycteridae | *Stauroglanis gouldingi* | 1191 | 697.9 | 8.1 | 201 | 2612 | 690 | 179 |
| Siluriformes | Trichomycteridae | *Trichomycterus aerolatus* | 1432 | 782.4 | 6.5 | 201 | 2245 | 818 | 233 |
| Tetraodontiformes | Tetraodontidae | *Takifugu rubripes* * | 991 | 196.0 | 1.8 | 96 | 740 | 184 | 0 |

The relationships we recover among the main lineages of Gymnotiformes (Figure 2) agree with previous studies that used mtDNA genomes (Elbassiouny *et al*., 2016) or exons (Arcila *et al*., 2017). Similar to the results in these studies, we resolve Apteronotidae, represented in our data set by *Sternarchorhamphus muelleri*, as the earliest diverging branch in the order. This placement of Apteronotidae disagrees with previous morphological and Sanger-based hypotheses which suggested either Gymnotidae (banded knifefishes of the genus *Gymnotus* and electric eel) (Tagliacollo *et al*., 2016) or only the electric eel *Electrophorus* (i.e., non-monophyletic Gymnotidae) (Janzen, 2016) were the sister group to all the other families.

**Figure 2.**
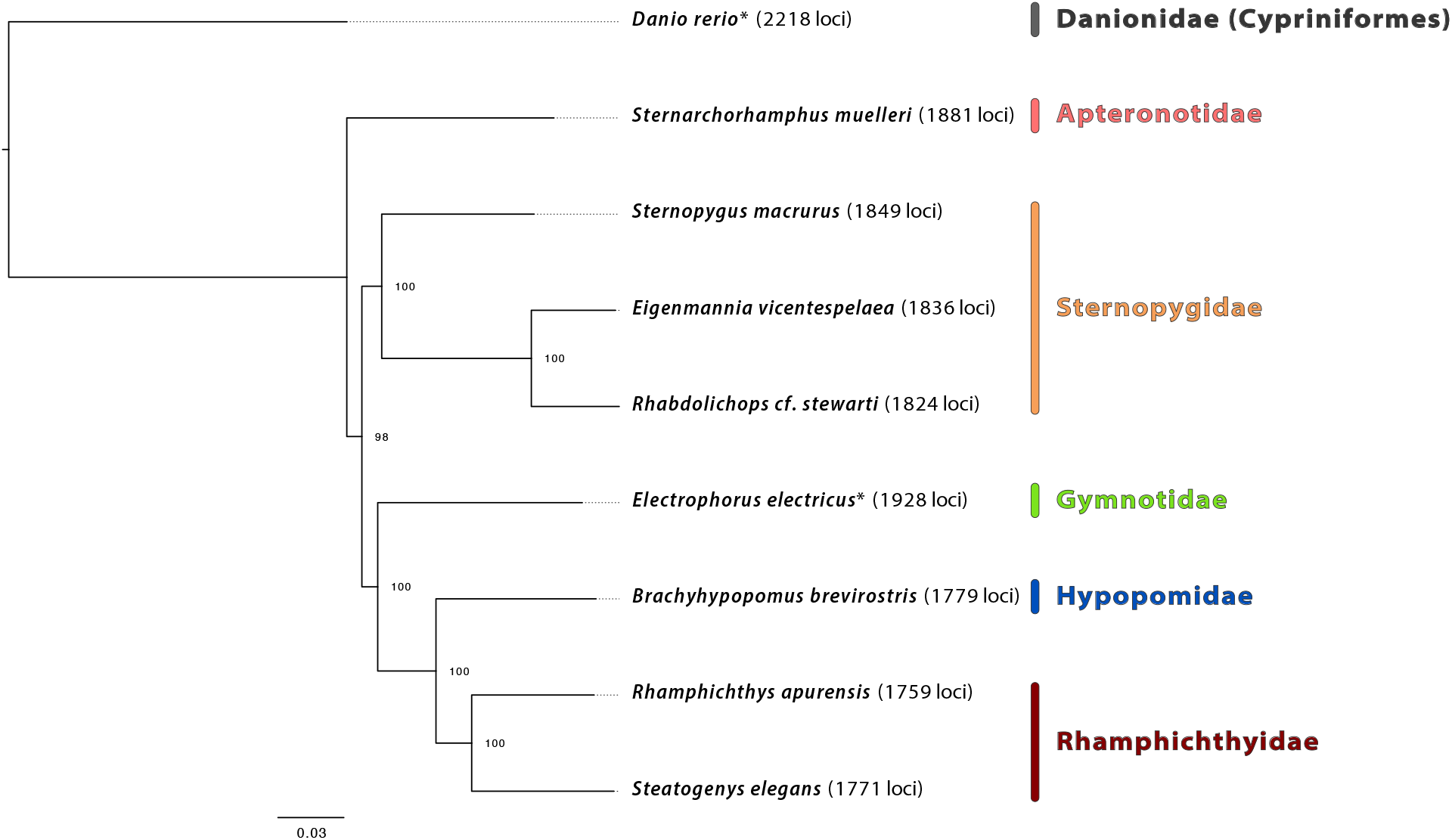
Maximum likelihood phylogenetic hypothesis of relationships among taxa comprising the Gymnotiformes data set with family names in color. *Danio rerio* is the outgroup taxon, and bootstrap support is indicated at each node. An asterisk by any taxon name indicates that these data were harvested, *in silico*, from existing genome assemblies, and the numbers in parentheses to the right of each taxon denote the count of loci enriched/harvested from that organism.

Our UCE results resolve representatives of the pulse-type families that produce electric organ discharges (Rhamphichthyidae (sand knifefishes) and Hypopomidae (bluntnose knifefishes)) as a monophyletic group while we resolved families producing electric signals in the form of waves (Apteronotidae (ghost knifefishes) and Sternopygidae (glass and rat-tail knifefishes)) as paraphyletic, a phylogenetic hypothesis that contrasts with previous studies that used morphology and Sanger sequencing data to suggest these families were monophyletic (Albert, 1998; Albert, 2001; Albert and Crampton, 2005; Janzen, 2016; Tagliacollo *et al*., 2016).

The differences we observed among the placement of gymnotiform families relative to previous studies reflects the confusing history of gymnotiform evolution where almost any possible hypothesis of relationships among gymnotiform families has been suggested (Triques, 1993; Gayet *et al*., 1994; Alves-Gomes *et al*., 1995; Albert, 1998; Albert, 2001; Albert and Crampton, 2005; Janzen, 2016; Tagliacollo *et al*., 2016; Arcila *et al*., 2017). These conflicts may arise from a very rapid diversification event that occurred around the origin of Gymnotiformes which created an evolutionary history muddled by incomplete lineage sorting. The causes of these incongruences and methods to increase consistency in the inferences drawn from UCE data are being explored as part of a separate study (Alda *et al*., In Review).

### Anostomoidea data set

The Anostomoidea data set (Table 1) was the second of two “young” ostariophysan subclades we studied (crown age falls within 76-51 MYA (Hughes *et al*., 2018)), and we enriched an average of 1,272 UCE loci from members of this group. These UCE loci averaged 493 bp in length and represented 1,987 of the 2,708 loci (73%) that we targeted (Table 4). Alignments of these loci contained an average of 9 taxa (range 3–15). After alignment trimming, the 75% matrix included 879 UCE loci containing an average of 13 taxa (range 11–15), having an average length of 487 bp, a total length of 428,381 characters, and an average of 68 parsimony informative sites per locus. RAxML bootstrap analyses required 50 iterations to reach the MRE stopping point.

Our ML analyses (Figure 3) recover a clear division between the omnivorous/herbivorous Anostomidae (headstanders) and a clade of three fully or partially detritivorous families (Chilodontidae, Curimatidae and Prochilodontidae), a result also found by earlier, Sanger-based analyses (Melo *et al*., 2014; Melo *et al*., 2016; Melo *et al*., 2018). Relationships within Anostomidae match the Sanger-based results of Ramirez at al. (2016) and differ from the morphology-based hypothesis of Sidlauskas and Vari (2008) in the placement of *Anostomus* as sister to *Leporellus* (rather than *Laemolyta*). Relationships within Curimatidae are fully congruent with Vari’s (1989) morphological hypothesis and a recent multilocus Sanger phylogeny (Melo *et al*., 2018).

**Figure 3.**
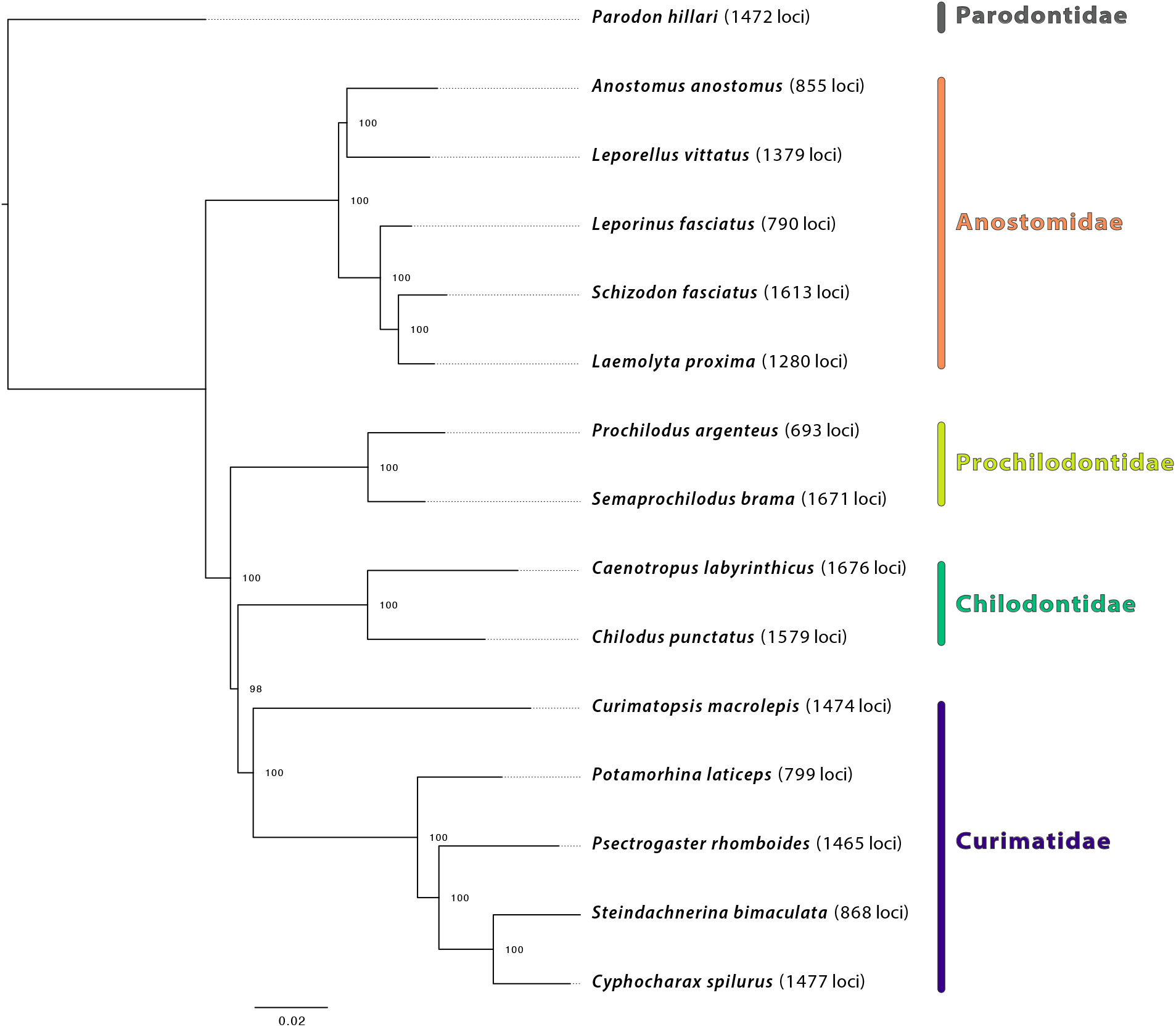
Maximum likelihood phylogenetic hypothesis of relationships among taxa comprising the Anostomoidea data set with family names in color. *Parodon hillari* is the outgroup taxon, and bootstrap support is indicated at each node. The numbers in parentheses to the right of each taxon denote the count of loci enriched from that organism.

We resolve Prochilodontidae and Chilodontidae as successive sister groups to Curimatidae. These results differ from recent Sanger sequencing studies (Oliveira *et al*., 2011; Melo *et al*., 2018) which reverse this order, and they also differ from Vari’s (1983) morphological hypotheses, which suggested Chilodontidae were sister to Anostomidae. Regardless of the exact relationships between Prochilodontidae, Chilodontidae, and Curimatidae, the resolution of branching order among these three primarily detritivorous characiform families is biologically interesting because either resolution implies a different and complex pattern of evolution in oral and pharyngeal dentition, the epibranchial organ, and numerous other anatomical systems. As noted for Gymnotiformes, the short branches associated with the near simultaneous origin of all three families may explain differences between this study and Sanger-based studies, and future work investigating these relationships would benefit from sampling more broadly across these families and more thorough phylogenetic analyses.

### Loricarioidei data set

The Loricarioidei data set (Table 1) represented an ostariophysan subclade of moderate age (crown age ~125 MYA (Rivera-Rivera and Montoya-Burgos, 2017)), and from these taxa we enriched an average of 1,379 UCE loci having an average length of 781 bp. Before trimming and matrix reduction, the alignments represented 2,176 of the 2,708 loci (80%) we targeted (Table 4), each alignment contained a mean of 9 taxa (range 3–15), and average alignment length was 566. After alignment trimming, the 75% matrix included 938 UCE loci containing an average of 13 taxa (range 11–15), having a total length of 608,044 characters, an average length of 648 bp, and an average of 261 parsimony informative sites per locus. RAxML bootstrap analyses required 50 iterations to reach the MRE stopping criterion.

The major relationships we resolve among families of Loricarioidei (Figure 4) are congruent with previous morphological hypotheses (Mo, 1991; Lundberg, 1993; Pinna, 1993; de Pinna, 1996; de Pinna, 1998), an earlier Sanger molecular hypothesis (Sullivan *et al*., 2006), and the exon-enrichment based molecular hypothesis of Arcila *et al.* (2017). Interestingly, we resolve family Scoloplacidae (spiny-dwarf catfishes) and family Astroblepidae (climbing catfishes) as successive sister groups to Loricariidae (armored catfishes), a placement reported by other studies (de Pinna, 1998; Sullivan *et al*., 2006) that suggests the loss of armor plating in Astroblepidae (de Pinna, 1998). Because relationships within this group remain controversial (Sullivan *et al*., 2006; Rivera-Rivera and Montoya-Burgos, 2017) and because Loricarioidei is the most diverse suborder of Neotropical catfishes (Sullivan *et al*., 2006), additional studies of interfamilial relationships and family status within the group are needed.

**Figure 4.**
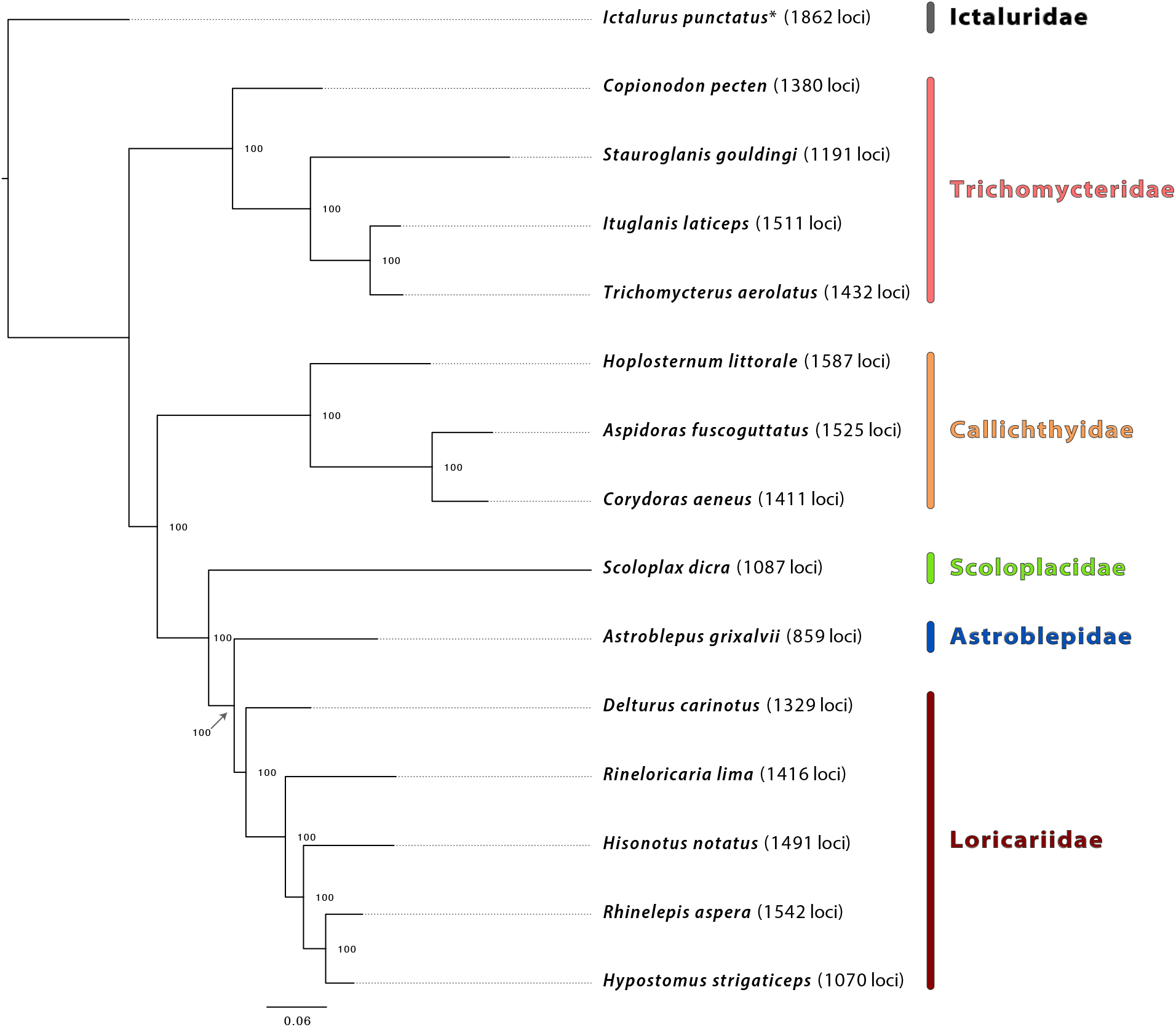
Maximum likelihood phylogenetic hypothesis of relationships among taxa comprising the Loricarioidei data set with family names in color. *Ictalurus punctatus* is the outgroup taxon, and bootstrap support is indicated at each node. An asterisk by any taxon name indicates that these data were harvested, *in silico*, from existing genome assemblies, and the numbers in parentheses to the right of each taxon denote the count of loci enriched/harvested from that organism.

### Characiformes data set

The Characiformes data set (Table 1) represented our second ostariophysan subclade of moderate age (~122 MYA (Hughes *et al*., 2018)), and we enriched an average of 1,701 UCE loci from each taxon having an average length of 784 bp (Table 4). Before trimming and matrix reduction, the alignments represented 2,493 of the 2,708 loci we targeted (92%), each alignment contained a mean of 15 taxa (range 3–22), and average alignment length was 526. After alignment trimming, the 75% data matrix included 1,399 UCE loci containing an average of 19 taxa (range 16–22), having a total length of 807,240 characters, an average length of 577 bp, and an average of 220 parsimony informative sites per locus. RAxML bootstrap analyses required 50 iterations to reach the MRE stopping criterion.

The overall pattern of relationships we resolved for Characiformes (Figure 5) is similar to those from multilocus Sanger sequencing (Oliveira *et al*., 2011) or exon-based (Arcila *et al*., 2017) studies. For example, our results include separation of the African Citharinoidei (Citharinidae and Distichodontidae) from other characiforms in the earliest divergence within the order and resolution of Crenuchidae (Neotropical darters) as sister to all other members of Characiformes (suborder Characoidei). Within Characoidei, we resolved two major lineages: one comprising Ctenoluciidae (pike-characins), Lebiasinidae (pencilfishes), Acestrorhynchidae (dogtooth characins), Bryconidae (dorados and allies), Triportheidae (elongate hatchetfishes) and members of the hyperdiverse family Characidae (tetras) and the other including a monophyletic superfamily, Anostomoidea (headstanders, toothless characiforms and relatives), that is closely aligned to Serrasalmidae (piranhas and pacus), Hemiodontidae (halftooths) and Parodontidae (scrapetooths), and more distantly related to Erythrinidae (trahiras) and the second clade of African families Alestidae and Hepsetidae. Within Characoidei, the short branches connecting internodes along the backbone of the phylogeny reflect previous results suggesting a rapid initial diversification of families within this suborder (Arcila *et al*., 2017; Chakrabarty *et al*., 2017).

**Figure 5.**
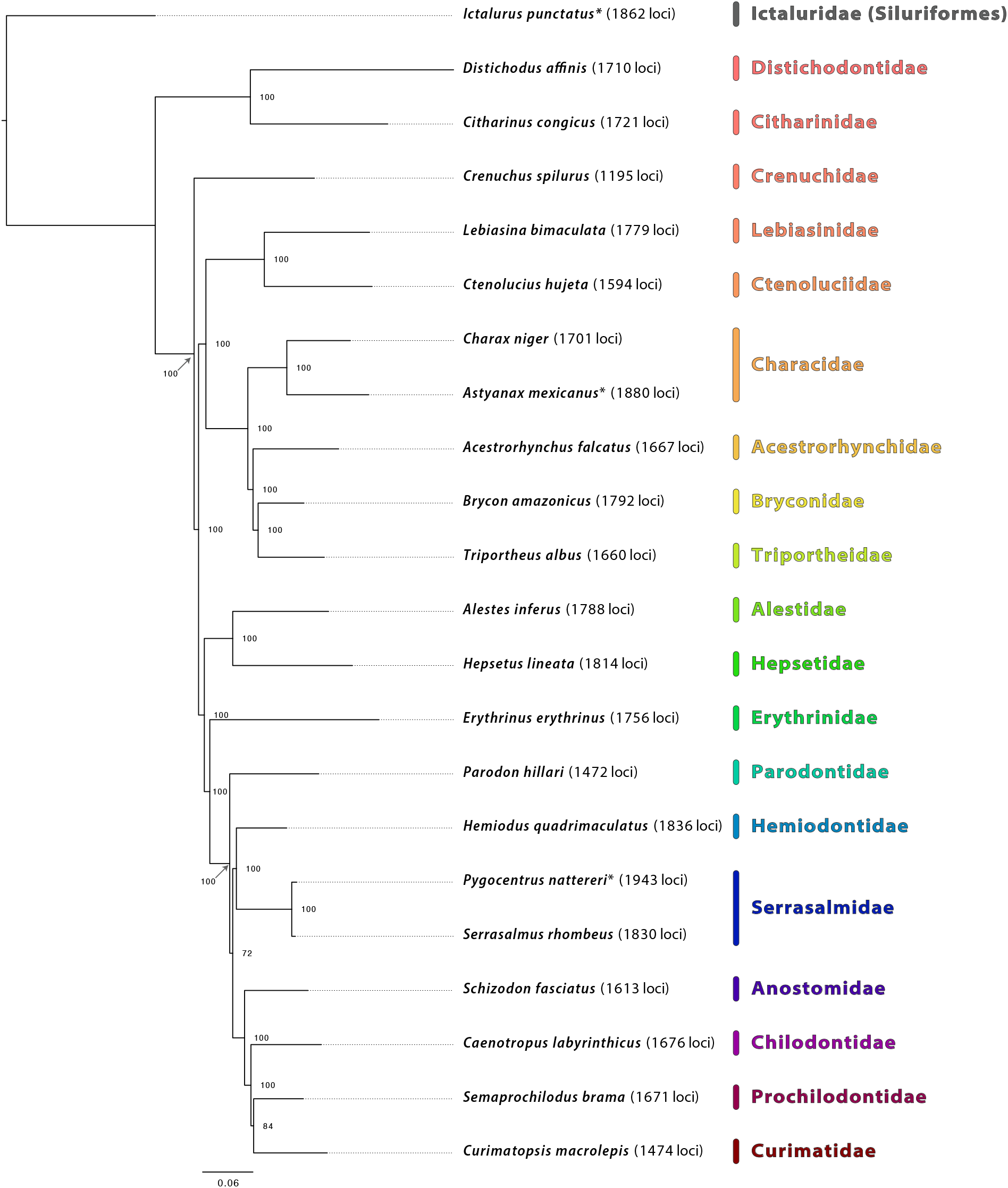
Maximum likelihood phylogenetic hypothesis of relationships among taxa comprising the Characiformes data set with family names in color. *Ictalurus punctatus* is the outgroup taxon, and bootstrap support is indicated at each node. An asterisk by any taxon name indicates that these data were harvested, *in silico*, from existing genome assemblies, and the numbers in parentheses to the right of each taxon denote the count of loci enriched/harvested from that organism.

### Otocephala data set

The Otocephala data set (Table 1) represented the oldest clade of fishes we investigated (~193 MYA (Hughes *et al*., 2018)), and we created this data set by combining enrichment data from select lineages used in the data sets above with enrichment data collected using the same array from taxa representing Clupeiformes and Cypriniformes (Table 1). To these empirical data, we integrated *in silico* data harvested from even more distant outgroups to show that the ostariophysan bait set is useful to study these other groups and also to demonstrate that it recovers reasonable relationships among these various lineages. From the taxa in this data set where we used targeted enrichment to collect data, we enriched an average of 1,447 UCE loci having an average length of 784 bp. When we combined these data with the *in silico* data, the alignments represented 2,573 of 2,708 loci (95%), each alignment contained a mean of 11 taxa (range 3–21), and average alignment length was 445 bp. After alignment trimming, the 75% data matrix included 658 UCE loci containing an average of 17 taxa (range 15–21), having an average length of 384 characters, a total length of 252,749 characters, and an average of 146 parsimony informative sites per locus. RAxML bootstrap analyses required 350 iterations to reach the MRE stopping criterion.

The branching order we resolve among Lepisosteiformes, Anguilliformes, Osteoglossiformes, and Euteleostei relative to the otocephalan ingroup (Figure 6) is similar to the pattern of major relationships among these fish groups resolved by other phylogenomic studies (Faircloth *et al*., 2013; Hughes *et al*., 2018). Similarly, the UCE data we enriched from lineages representing Clupeiformes and Cypriniformes produced the same phylogenetic hypothesis for the branching order of these groups relative to the Characiphysi (Characiformes + Gymnotiformes + Siluriformes) as seen in other genome-scale (Hughes *et al*., 2018) and Sanger sequencing (Near *et al*., 2012; Betancur-R *et al*., 2013) studies. Relationships among the orders comprising otophysans are similar to some genome-scale studies and different from others, reflecting the difficulties noted when studying these groups (reviewed in Chakrabarty *et al*., 2017; Arcila *et al*., 2017).

**Figure 6.**
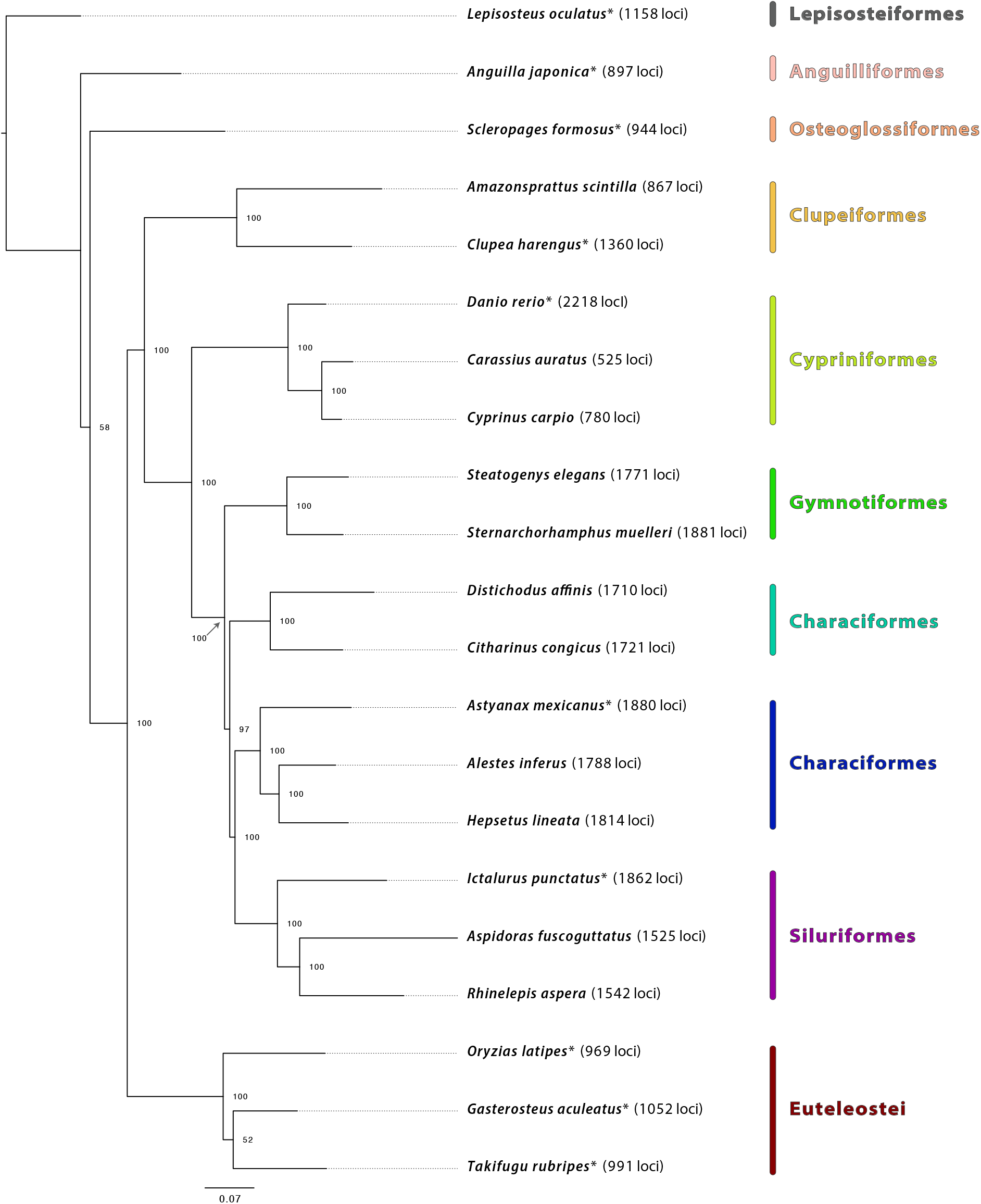
Maximum likelihood phylogenetic hypothesis of relationships among taxa comprising the Otocephala data set with family names in color. *Lepisosteus oculatus* is the outgroup taxon, and bootstrap support is indicated at each node. An asterisk by any taxon name indicates that these data were harvested, *in silico*, from existing genome assemblies, and the numbers in parentheses to the right of each taxon denote the count of loci enriched/harvested from that organism.

## DISCUSSION

The bait set that we designed effectively collected data from the majority of the 2,708 UCE loci that we targeted across the four ostariophysan subclades we investigated: averaging across all of our experiments except the Otocephala data set, which included many genome-enabled taxa, we enriched an average of 2,229 of the 2,708 loci (82%). This bait set also performed well when enriching putatively orthologous loci from *Amazonsprattus scintilla* (Clupeiformes). Because of our success enriching loci from Clupeiformes, which are a close outgroup to Ostariophysi, and despite our lack of a lineage representing Gonorynchiformes, we refer to this bait set as targeting Ostariophysi/ostariophysans rather than smaller subclades within this group.

As detailed above, the data we collected using the ostariophysan bait set reconstruct reasonable phylogenetic hypotheses for all datasets, despite low taxon sampling (less than 1% of diversity for the overall study and less than 5% in Anostomoidea, the most densely sampled subclade). By reasonable, we mean that the phylogenetic hypotheses we resolved largely agree with previous investigations using multilocus Sanger sequencing data or genome-scale data collection approaches. Where we observed differences from some prior studies were those relationships having very short internal branches suggesting rapid or explosive radiation of a particular clade. These areas of treespace are hard to reconstruct (Pamilo and Nei, 1988; Maddison, 1997; Maddison and Knowles, 2006; Oliver, 2013), and many current studies are focused on analytical approaches that produce the most accurate phylogenetic hypothesis given the data. The congruence of our results with stable parts of the trees inferred during these earlier studies and the overall ability of this bait set to pull down significant proportions of the targeted loci suggests that our ostariophysan bait set provides one mechanism to begin large-scale data collection from and inference of the relationships among the more than 10,000 species that comprise Ostariophysi, many of which have never been placed on a phylogeny.

Future work should explicitly test the effectiveness of this ostariophysan bait set for enriching loci from Gonorynchiformes, the smallest ostariophysan order and a group for which tissue samples are few. Similarly, this bait set should be tested in Alepocephaliformes, an enigmatic order of marine fishes that may form a close outgroup to Ostariophysi. Despite those gaps, our *in silico* results suggest: (1) that this bait set may be useful in even more distant groups like Osteoglossiformes or Euteleostei, and (2) the exciting possibility that we may be able to create a large (>1000–2000 loci), combined bait set targeting orthologous, conserved loci that are shared among actinopterygians to reconstruct a Tree of Life spanning the largest vertebrate radiation.

## ACKNOWLEDGEMENTS

We thank the curators, staff, and field collectors at the institutions listed in Table 1 for loans of tissue samples used in this project. This work was supported by grants from NSF to BCF (DEB-1242267), BLS (DEB-1257898), and PC (DEB-1354149) and FAPESP to CO (14/26508-3), BFM (16/11313-8), FFR (14/05051-5), and LEO (14/06853-8). Animal tissues collected as part of this work followed protocols approved by the University of California Los Angeles Institutional Animal Care and Use Committee (Approval 2008-176-21). Portions of this research were conducted with high-performance computing resources provided by Louisiana State University (http://www.hpc.lsu.edu).

## DATA ACCESSIBILITY

Sequence data from *A. albifrons* and *C. paleatus* used for locus identification are available from NCBI BioProject PRJNA493643, and sequence data from enriched libraries using the ostariophysan bait set are available from NCBI BioProject PRJNA492882. The ostariophysan bait design file is available from FigShare (doi:10.6084/m9.figshare.7144199), where it can be updated, if needed. A static copy of the bait design file and all other associated files, including contig assemblies, UCE loci, and inferred phylogenies are available from Zenodo.org (doi:10.5281/zenodo.1442082).

